# Deviance detection via competitive inhibition between local neocortical ensembles

**DOI:** 10.1101/2025.01.08.631952

**Authors:** Ryan V. Thorpe, Christopher I. Moore, Stephanie R. Jones

## Abstract

The process by which neocortical neurons and circuits amplify their response to an unexpected change in stimulus, typically referred to as deviance detection (DD), has traditionally been thought to be the product of specialized cell types and/or routing from distinct brain areas. Here, we explore a different theory, whereby DD emerges intrinsically from local network-level interactions within a neocortical column. We propose that deviance-driven neural dynamics are generated by ensembles of excitatory and inhibitory neurons that have a fundamental inhibitory connectivity motif: competitive inhibition between reciprocally connected neural representations under modulation from feed-forward selective (dis)inhibition. Implementing this motif in two computational models with different levels of biophysical abstraction, we were able to simulate a variety of phenomena pertaining to the experimentally observed shifts in neural tuning during DD across neurons, time, and stimulus history. We further tested hypotheses related to our theory and examined the robustness of emergent phenomena consistent with prior experimental observations. Our results show that ensemble priming via competitive inhibition under modulation from selective (dis)inhibition can serve as a local mechanism for encoding short-term stimulus memory, enabling deviance-driven shifts in stimulus representation. This work establishes a novel theoretical paradigm that resolves previously confounding aspects of predictive sensory processing in Neocortex, and we provide a number of corollary predictions that can be tested in future in vivo studies.

## Introduction

Real world behavior occurs in real time, with threats and opportunities constantly, often rapidly, evolving. Detecting subtle changes in ongoing input is essential to avoiding an approaching predator or noticing key resources while scanning an environment. Accordingly, ‘deviants’ in ongoing perceptual information often powerfully enhance or redirect salience, increasing the likelihood of detecting even subliminal stimuli [1–3].

This enhanced perceptual sensitivity is matched by neocortical responses that show distinct sensitivity to unexpected deviant stimuli, an amplification mechanism widely regarded as crucial for sensory processing [4–10]. Among other hallmark phenomena, groups of neurons in Sensory Neocortex exhibit deviance detection (DD) by increasing their average action potential spike rates in response to deviant stimuli [11–13]. Decreases in firing in specific subsets of cells and shifts in macroscopic features of electroencephalography (EEG), magnetoencephalography (MEG), and local field potential (LFP) are also commonly observed [12, 14–24]. How specific neurons, comprising functional ensembles, shift their stimulus-evoked response profiles on deviant trials remains an open-ended question.

Two classes of theories, both built on widely accepted neural mechanisms, have been offered to explain the observed increase in deviance-driven neural responses. In the ‘bottom-up’ account, a local increase in neocortical firing occurs when a change in sensory input recruits a new (i.e., unadapted) feed-forward pool of neurons [1, 4, 24]. For example, when a visual or somatotopic topographic position is repeatedly stimulated and then a new position is stimulated, activation of neurons along a novel afferent pathway should provide unadapted, and therefore more robust, responses in the primary sensory neocortical representation [13, 25]. In the ‘top-down’ account, inhibition evoked by feedback from higher-order neocortical areas suppresses the representation of predicted stimulus inputs. When a novel variation is presented—even one that activates pyramidal neurons in the same topographic neocortical column as the predicted stimulus—this inhibition is avoided, leading to an enhanced response [4, 5, 8, 11, 12]. One example cited for such a mechanism is that an unexpected change in the navigation-dependent representation of an expected feature of the visual scene can drive a deviance response in Primary Visual Neocortex (V1) [26].

Both models likely contribute to augmented neural responses [8, 9, 13], granting the Neocortex multiple mechanisms for calculating and propagating a prediction error signal it can use to guide learning and behavior [2, 4, 5]. However, these models have limitations. Both circuit mechanisms require a shift in the stimulus-activated afferent pathway, while amplified spike rate responses occur even with deviant stimuli that are not expected to recruit a new group of differently tuned neurons. For instance, DD manifests in response to a sudden decrement in the intensity of an otherwise identical stimulus [2, 19–21]. Further, DD emerges across neocortical areas: To our knowledge, every area thus far tested has revealed a form of this phenomenon [8, 9, 13, 23–25]. As such, a hierarchical model of predictive processing struggles to explain DD in frontal neocortical areas that ostensibly lack a ‘higher’ area to draw predictive input from.

To complement existing models and extend mechanistic understanding of DD, here we present a new framework, one that accounts for deviance-driven shifts in neural tuning and evoked spike rates and achieves DD through local dynamics within a single neocortical column. Our model explains how deviance-driven shifts in neural responses can emerge without targeting a new neural population specifically tuned to the deviant stimulus.

This model emerged in part to account for the findings of Voigts *et al*. [2], who conducted extensive neural recordings in mouse Primary Somatosensory Neocortex (SI) during the presentation of amplitude deviants, where the intensity of a vibrissa deflection, and ostensibly the magnitude of downstream recruitment of a common pool of neurons, changed on a deviant trial. Deviance-driven responses were layer-specific and particularly prominent at low stimulus magnitudes near perceptual threshold (i.e., within the dynamic range of an animal’s psychometric response; Figure 1a). Excitatory neurons in layer 4 (L4) exhibited a supralinear spike rate response to an increase in stimulus intensity, a deviance-driven response that exceeded the change anticipated by standard intensity coding accounting for synaptic adaptation. In parallel, a decrease in stimulus intensity caused a sublinear response (Figure 1b). In sum, deviant stimuli evoked responses with exaggerated, and putatively enhanced, encoding of the direction of intensity change. In contrast to the tuning observed in L4, L2/3 neurons showed more complex tuning. In addition to subsets of cells that showed L4-like responses, other L2/3 neurons showed *increased* firing rates to a deviant *decrease* in stimulus amplitude, and conversely, decreased firing to an increase in stimulus amplitude (Figure 1c). The emergence of such tuning in response to subtle changes in the intensity of tactile stimuli is consistent with other findings showing high sensitivity to complex features and deviations, such as the sensitivity of L2/3 neurons to navigational expectations and/or sensorimotor context across sensory systems [11, 26–32]. As such, we refer to the distinct response properties of L2/3 neural populations as ‘deviance-driven complex tuning’. Deviance-driven complex tuning can emerge with a few variants of underlying ensemble dynamics (Figure 1d) that we discuss in detail below (see Results).

**Figure 1.**
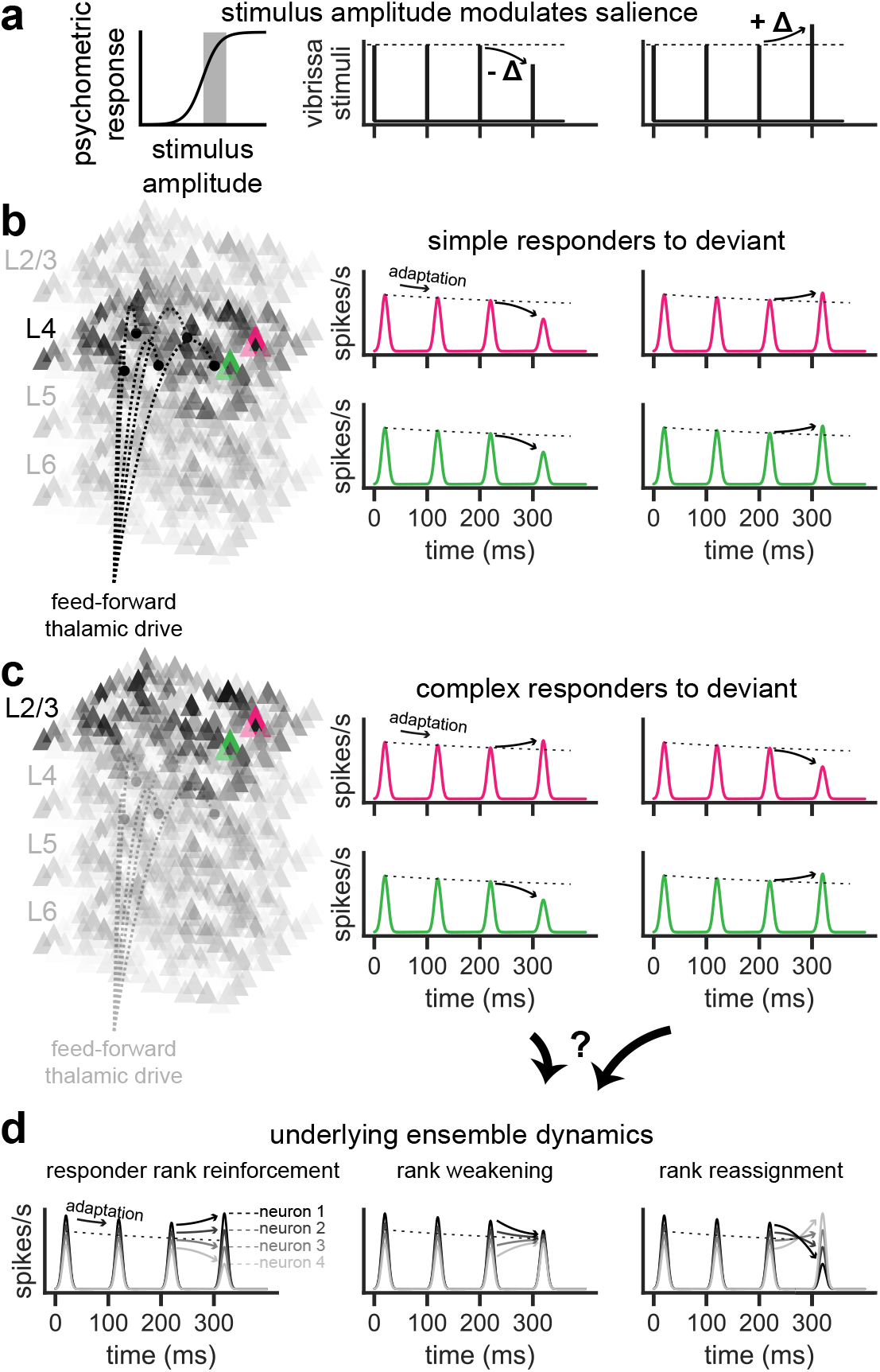
Complex tuning emerges in L2/3 with a deviant change in stimulus intensity. (a) Experimental stimulation paradigm, where vibrissa stimuli are presented that do not change canonical features of the sensory input (e.g., the direction of motion or topographic position) but rather the intensity of the stimulus. In Voigts *et al*. [2], such stimuli were applied within the dynamic range of the psychometric response. (b) Depictions of the single-unit spike rate evoked responses observed in L4 of mouse barrel Neocortex for two excitatory neurons (pink and green). Excitatory neurons were found to show a supralinear response to a deviant increase or a sublinear response to a deviant decrease in intensity of sensory drive [2]. (c) In L2/3, a more heterogeneous and complex set of tuning properties emerged. Some neurons showed L4-like responses, while others produced paradoxical increases in firing in response to deviant decreases in amplitude, or vice versa [2]. (d) Three hypothetical scenarios that could capture the ensemble dynamics underlying the transient shift in deviant tuning profiles of complex responders in L2/3.

Key features of the response properties in L2/3 provide constraints on possible mechanistic (model) implementations. First, the deviance-driven complex tuning observed in Voigts *et al*. [2] emerged rapidly at latencies < 100 ms. Second, subtle optogenetic manipulation of L6 firing—firing that did not cause a detectable change in the overall firing of L6 but simply shifted the tuning of individual neurons in that layer—changed L2/3 deviance tuning and animal behavior. L6 manipulation removed deviance-driven complex tuning in L2/3, shifting responses to express those seen in L4 [2]. Further, this same modest intervention removed the behavioral benefit of deviance for stimulus detection [2]. These results suggest that deviance-driven complex tuning in L2/3, and the putative benefit of this encoding, emerges at least in part through rapid intracolumnar circuit mechanisms.

Several lines of evidence suggest that local interactions between pools of inhibitory interneurons could be crucial to deviance-driven complex tuning in L2/3. Physiological studies have shown functional interneuron-interneuron inhibitory interconnectivity in specific layers of Neocortex [33–36]. The detailed high-resolution anatomical mapping of putative synaptic interconnectivity reinforces these findings, showing robust connections between distinct inhibitory interneuron motifs in the upper layers of V1 [37]. Perhaps most importantly, recent calcium imaging studies by Deister *et al*. [38] in L2/3 of SI and V1 found that two distinct pools of fast-spiking, parvalbumin-positive (PV) interneurons predicted detection success for tactile or visual stimuli.

These two interneuron pools showed opposed firing rate changes, rate decreases or increases, respectively. Modeled interneuron interactions demonstrated that cross-inhibitory suppression between these pools could account for key phenomena that also predicted detection success, such as increased correlation and spiking in a discrete excitatory neuron ensemble, and decorrelation of the remaining, non-participating cells [38]. In this motif, excitatory neurons are clustered according to stimulus preference (including tuning for task success) [39–42] and distinct pools of interneurons that target pyramidal somata also inhibit each other. This opponency allows for selective (dis)inhibition of specific, task-predictive ensembles. Further, this motif is particularly powerful when recurrent inhibition leads to competition between neural representations and can also be modulated by external ‘top-down’ and/or ‘bottom-up’ input targeting either representation [38, 43–46].

We sought to determine if such an inhibitory motif could act as a local, circuit-driven mechanism for the emergence of deviance-driven complex tuning. Further, we sought to account for experimentally observed shifts in feature discrimination [47–49] concurrent with the adaptation of neural tuning profiles (i.e., sharpening and/or broadening) [49–52]. We used simulations from two computational models of different scales to test if and how distinct network configurations leverage inhibition to prime and shape the response of neuron ensembles in service of complex tuning, and more broadly, DD. First, we used a discrete Wilson-Cowan neural mass model to test generalized principles of inhibitory dynamics between two arbitrary competing neural representations [53, 54]. Our results demonstrate that reciprocal inhibition between at least two competing ensembles of neurons (competitive inhibition; CI) under modulation from selective (dis)inhibition (SDI) explain the emergence of deviance-driven complex tuning. Second, to show how this mechanism can be implemented under more realistic conditions, we extended the framework to a biophysically-detailed neocortical column model constrained by laminar structure, anatomical connectivity, and distinct neuron types with fast and slow excitation and inhibition [55]. With a number of corollary predictions that can be tested in future in vivo studies, we show that a simple network connectivity motif promotes deviance-driven complex tuning, short-term sensory context storage (i.e., stimulus memory), and that DD does not require specialized input from other brain areas—a novel theoretical paradigm that resolves previously confounding aspects of sensory encoding and predictive processing in Neocortex.

## Results

### Deviance-driven complex tuning emerges with inhibitory competition and selective (dis)inhibition

We began by proposing that only a handful of network dynamic scenarios could possibly serve as a basis for deviance-driven complex tuning at the neural ensemble level (Figure 1d). Namely, neurons tuned to a stimulus can respond to deviant increases or decreases in stimulus intensity with a more pronounced ordinal position (responder rank reinforcement; Figure 1d, left panel) or diminished ordinal position (responder rank weakening; Figure 1d, middle panel) relative to other stimulus-responsive units ranked by evoked spike rate magnitude. If a sharp tuning profile can give way to a weakened tuning profile, the ordinal position of various stimulus-responsive units can, in theory, invert. Complete inversion of the neurons’ ordinal position compared to baseline would establish a sharp, yet reversed, tuning profile (responder rank reassignment; Figure 1d, right panel). We emphasize that in each of these cases, a deviant decrease in stimulus amplitude does not necessarily decrease the mean evoked response across neurons. Instead, it selectively amplifies the response of some neurons at the expense of others in a manner that maintains, if not increases, the net salience of the neocortical circuit’s output. Each ensemble-level response intrinsically supports divisive normalization as a foundational computational principle—a network constraint we explore below from a dynamical systems perspective [56, 57].

The dynamics of responder rank reinforcement, rank weakening, and/or rank reassignment represent cases of transient shifts in stimulus preference. In order for such shifts to occur from mechanisms intrinsic to a local network in Neocortex, we assumed that neural tuning within a local population is determined by two factors: preference from feed-forward anatomical connections and priming from local recurrent interactions [43, 46, 58]. We therefore introduced a motif where neural ensembles are governed by the processes of CI and SDI, described as follows.

Consider two ensembles of interconnected neurons—each encoding a distinct feature or combination of features of the environment through the mean spike rate response across constituent neurons—that naturally receive different levels of afferent excitation evoked by an external stimulus. If these ensembles (or neural ‘representations’) exhibit uniform, non-clustered connectivity, differences in evoked activity readily propagates from one ensemble to the next and equilibrates quickly due to mutual excitation (ME). If, on the other hand, the two ensembles exhibit a high degree of clustered connectivity and reciprocally inhibit each other (Figure 2a), evoked activity produces CI that augments the difference between each ensembles’ response. With multiple repetitions of the same stimulus, CI drives distinct ensemble responses apart, aiding in feature discrimination that adapts with each repetition (i.e., adaptive sharpening of the neural tuning; Figure 2b; see also Machens *et al*. [45] for an example applying CI in a model for stimulus interval discrimination). The neural response across neurons intrinsically avoids saturation, and the equilibrium state becomes markedly time and repetition-dependent as governed by the shifting excitation/inhibition (E/I) balance *between* ensembles [59]. In this latter case, the neuronal representation across ensembles self-normalizes and from the perspective of a naive observer, appears to shift over time and space. However, if evoked (dis)inhibition via a parallel feed-forward pathway (e.g., from another pair of ensembles or local neocortical layer) selectively dampens the activity of one representation more than the other in a stimulus-dependent manner (termed here as the process of selective [dis]inhibition, SDI), the diverging effects of CI could be modulated to produce apparent shifts from the ensembles’ baseline tuning (Figure 2c).

**Figure 2.**
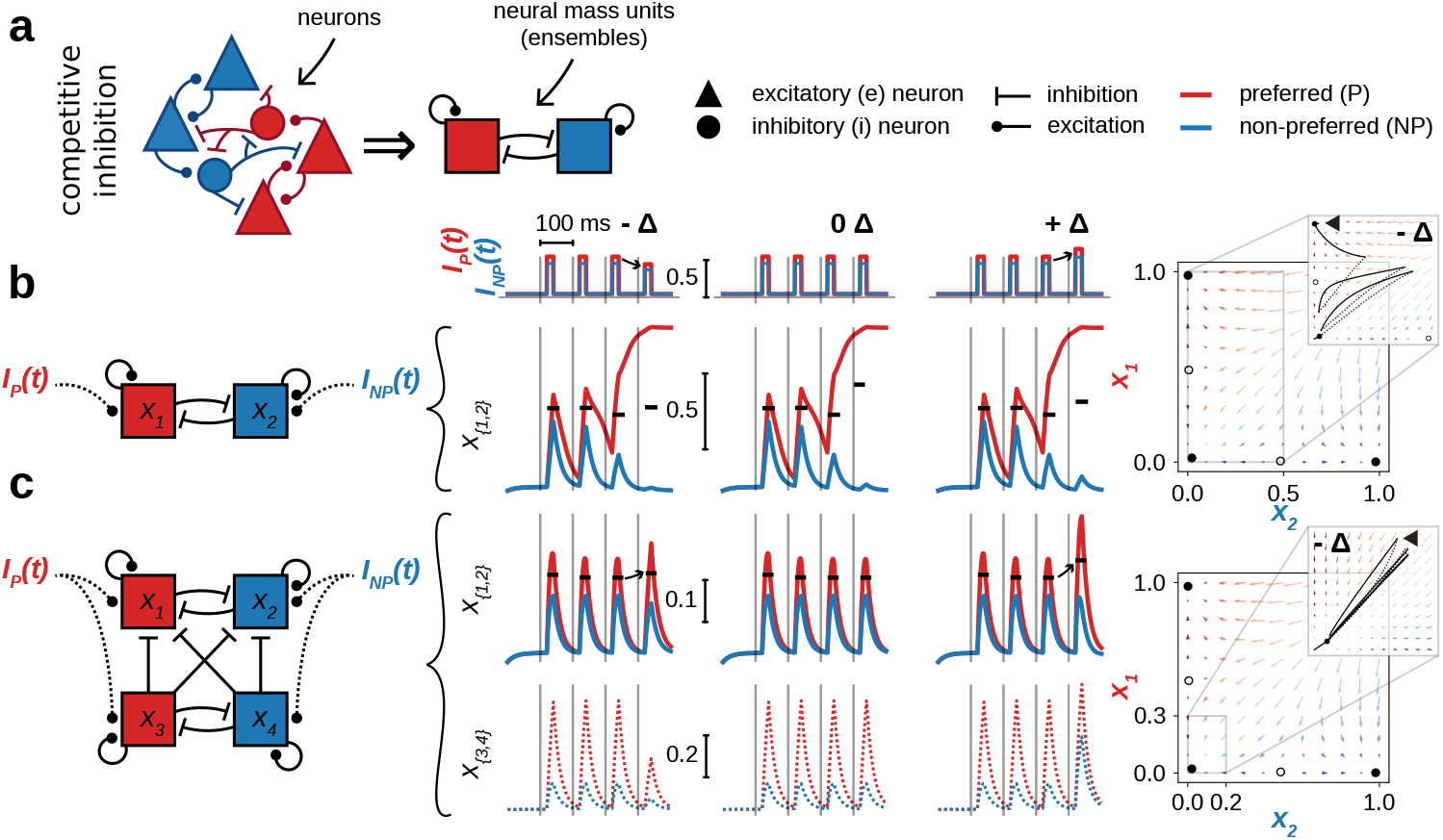
Competitive inhibition (CI) between neural representations modulated by selective (dis)inhibition (SDI) from another functional neural layer facilitates net amplification of the deviance-driven response and deviance-driven complex tuning. (a) CI requires at least two neuron ensemble representations that reciprocally inhibit each other yet allow excitation to spread among constituent neurons of each respective ensemble. One ensemble is maximally tuned to an arbitrary stimulus (preferred, red) while the other is sub-optimally tuned to the same stimulus (non-preferred, blue). (b) Simulation of diverging responses to repetitive sensory afferent drive within a two-dimensional model of CI. Simulations comprise a sequence of representation-specific input (*I*_*P*_ (*t*) and *I*_*NP*_ (*t*); top) producing standard stimulus-evoked responses followed by a negative deviant (middle-left), no deviant (middle-center), or positive (middle-right) deviant response. On the far right, the resting-state phase plane with stable and unstable fixed points (solid versus hollow dots, respectively) contains competing basins of attraction that a given simulation trajectory (inset) navigates between following each stimulus perturbation. (c) Same as in (b), except with a 4-dimensional network architecture that facilitates a deviance-driven mean spike rate increase and complex tuning in the upper units, *x*_1_ and *x*_2_. The connectivity motif enforces CI in two functional layers (upper and lower) and uni-directional cross-laminar inhibition from the lower to upper units. Evoked activity in the lower layer produces SDI of evoked activity in the upper layer. For both negative and positive amplitude deviants, the model responds with a non-decreasing average response across upper units (black tick marks).

We hypothesized that CI between at least two ensembles of neurons modulated by SDI from another functional layer (Figure 2c) can account for one or more of the ensemble dynamics scenarios subserving deviance-driven complex tuning (Figure 1d). Importantly, such a system would need to produce complex tuning while also maintaining, if not increasing, the total magnitude of the evoked output of the network, a hallmark characteristic of DD [11–13]. We first explored this idea in a discrete (space-clamped) Wilson-Cowan model of neural mass firing rates given by

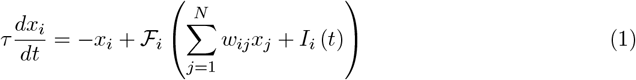

where ℱ_*i*_ is a sigmoid activation function for the *i*^*th*^ unit (ensemble) that increases or decreases with endogenous recurrent and exogenous input *w*_*ij*_*x*_*j*_ and *I*_*i*_(*t*), respectively [53, 54]. To test that the theoretic mechanisms of CI+SDI are (at minimum) sufficient for producing deviance-driven complex tuning in a manner that maintains, if not increases, the total magnitude of the network’s evoked output, we used the system defined in Equation 1 with *N* = 4 to establish a fundamental 4-dimensional network motif that reduces our hypothesized mechanisms down to their simplest form (Figure 2c; note that CI without SDI is shown for illustrative purposes in Figure 2b). As explored later in this study, the constraints on this motif can be loosened to represent general inhibitory ensemble interactions without CI and/or without SDI.

In the neural mass model defined in Equation 1, *x*_*i*_, can be thought of as the proportion of constituent neurons of the *i*^*th*^ ensemble that are firing at a given point in time. Each neural mass unit (ensemble) comprises a distinct neural representation that is either preferred (red) or non-preferred (blue) by exogenous stimulus-evoked drive. Exogenous excitatory drive is injected into the network through *I*(*t*), a time-dependent step function designed to emulate the stimulus-evoked afferent input entering each ensemble of the neocortical column. As shown at the top of Figure 2b, *I*(*t*) is slightly larger for red than blue (representing greater evoked drive targeting the preferred ensemble) when it is in its high state beginning at 20 ms post-stimulus, and then returns to a common low state at 40 ms post-stimulus in a sequence of four stimuli. On the fourth stimulus repetition, the magnitude decreases or increases across representations by a scalar factor corresponding to a deviant decrease in stimulus strength. Note that the relative balance of afferent input for the preferred compared to non-preferred stimulus (*I*_*P*_: *I*_*NP*_) across standard and deviant trials remained constant: only the mean peak value across *I*_*P*_ and *I*_*NP*_ changed.

In addition to exogenous drive, the *i*^*th*^ neural mass unit also receives input through recurrent connections with itself and other neighboring units, as determined by connectivity weights *w*_*ij*_ from the *j*^*th*^ unit (Equation 1). The CI (Figure 2b) and CI+SDI network (Figure 2c) motifs imply a few fundamental connectivity constraints. Connection weights constrained to be positive represent excitatory connections that propagate excitation throughout constituent neurons of an ensemble, whereas connection weights constrained to be negative represent inhibitory connections in service of CI between ensembles in a layer and/or SDI between layers.

CI promotes diverging neural representations that augment with each presentation of stimulus-driven afferent drive (Figure 2b). Specifically, we see a diverging sequence of neural responses between two competing neural mass units (red versus blue) in response to a sequence of repetitive of drive, regardless of whether or not the final repetition in the sequence is deviant (Figure 2b, middle-left through middle-right). On the final stimulus, the dynamic state of the network has fallen into the upper attractor basin favoring the preferred representation, where it produces tonic spiking due to runaway excitation and minimal activity in the non-preferred representation.

With the addition of a modulatory layer that provides SDI via uni-directional cross-laminar inhibition to the same units depicted in Figure 2c, the diverging effect of CI in the upper layer is dampened. When SDI is decreased or increased based on the direction of the deviant (Figure 2c, middle-left and middle-right), the equilibrium of the network shifts, and the mean evoked response across upper layer units (black tick marks) not only refrains from decreasing on deviant trials, but increases for both negative (2c, middle-left) and positive deviants (2c, middle-right). Further, the network exhibits an increase in the difference between the red and blue evoked responses for both types of deviants compared to baseline, with red dominating. This response is an example of responder rank reinforcement as introduced in Figure 1d.

To verify that the CI+SDI motif (Figure 2c) is capable of responding uniquely to deviant stimuli that swap which feature is preferred instead of decreasing the total intensity across all features, we modified network connectivity to either produce slowly converging or diverging evoked responses, and then simulated a feature-swap deviant (Figure S1). Given that such a scenario maintains a constant mean level of evoked afferent drive even for deviant stimuli, we expected it to amplify the history-dependent nature of the CI+SDI motif so that the magnitude of the average deviance-driven response across representations increases depending on the number of prior stimuli of a particular feature preference. Indeed, this is exactly what we observed: for deviants that succeed 1, 2, 3, or 4 prior repetitive stimuli (Figure S1a,c), the magnitude of the average deviant response gets progressively larger (Figure S1b,d).

While other forms of deviant stimuli and deviance-driven complex tuning can emerge with a change in network parameters, this example demonstrates the overarching utility of CI and SI: a local neocortical circuit (or other brain area for that matter) can maintain, if not amplify, the total deviance-driven response through a common connectivity motif responsible for deviance-driven complex tuning. The network motif is uniquely designed to selectively amplify the response of a specific neural subpopulation at the expense of another, which is a process that can shift dynamically in response to a deviant change in magnitude or feature preference in service of DD.

### Inhibitory competition primes the network by facilitating adapting neural representations

We then sought to determine if and how CI and SDI interact to support both adaptation and deviance-driven shifts in neural tuning. CI intrinsically augments the difference between competing ensemble representations, priming the network for a deviant response with each repetition of a stimulus. Yet, as demonstrated in Figure 2c, the effects of CI can be dampened or enhanced by SDI based on the level of stimulus-evoked afferent drive. Numerous prior studies report that the stimulus tuning of neural representations shift over time, particularly when a given stimulus has been presented repetitively or sampled frequently by the peripheral nervous system within a short time window [49, 50, 52, 60]. It has been suggested that adaptive sharpening (i.e., the narrowing of a neuron’s tuning to external stimuli, similar to the ensemble-level phenomena of responder rank reinforcement) subserves the psychophysical phenomenon of enhanced feature discrimination due to stimulus repetition [47, 48, 51, 52, 60–64]. While many cellular and circuit mechanisms including short-term synaptic depression are likely involved in facilitating stimulus-specific adaptation [49, 51, 52, 64–68], here we assumed that local inhibition plays a prominent role in determining the responsiveness of latent ensembles of neurons [27, 32, 69].

We specifically asked if the CI+SDI network motif has a parameter regime where adaptive sharpening of neural tuning and deviance-driven complex tuning emerge, all while producing an average deviance-driven non-decreasing (DnD) response. It could be that deviance-driven complex tuning only emerges within a specific regime where adaptive sharpening cannot. Or, the average deviance-driven response might always decrease on negative deviant trials. Either way, a null result (where less than all desired phenomena are produced by the network) would invalid the CI+SDI motif as a possible network architecture underlying DD.

To test if the CI+SDI motif has a parameter regime where adaptive sharpening of neural tuning, deviance-driven complex tuning, and a DnD response all emerge, we estimated the likelihood of observing specific emergent phenomena given a uniform parameter prior distribution. Connectivity of the 4-dimensional CI+SDI motif is determined by six parameters, so we sampled 1 × 10^6^ random parameter values, each a 6-dimensional vector, to estimate the robustness of emergent dynamics across the parameter-space.

To control for the possibility that our desired emergent phenomena (adaptive sharpening of neural tuning, deviance-driven complex tuning, and a DnD response) might exist in a parameterspace outside the fundamental constraints of CI and SDI, we loosened the CI and SDI constraints by allowing the cross-representation connections to take a positive (excitatory) value, and cross-laminar inhibitory connections to take a maximal value of 0. Thus, a randomly sampled parameter could produce a network with ME between ensemble representations and/or silenced cross-laminar SDI.

We found that the network could indeed produce all three emergent phenomena (adaptive sharpening, deviance-driven complex tuning, and a DnD response); however, concurrence of such phenomena was rare and highly sensitive to network parameters. After discarding parameter configurations where the network produced a self-sustaining spike rate response through runaway excitation (termed here as ‘unstable’; 76.2%) and those that were stable yet produced adaptive broadening (12.1%), only 11.7% of the total 10^6^ samples produced adaptive sharpening (Figure 3). Of those, an even smaller percentage produced DnD responses (∼ 0.2% of the total original number of samples; Figure 3). Each parameter configuration that produced DnD responses also produced deviance-driven complex tuning, which we further categorized as either responder rank reinforcement, weakening, or reassignment for negative and positive deviants as defined in Figure 1d (Figure 3, bar-plot on the right).

**Figure 3.**
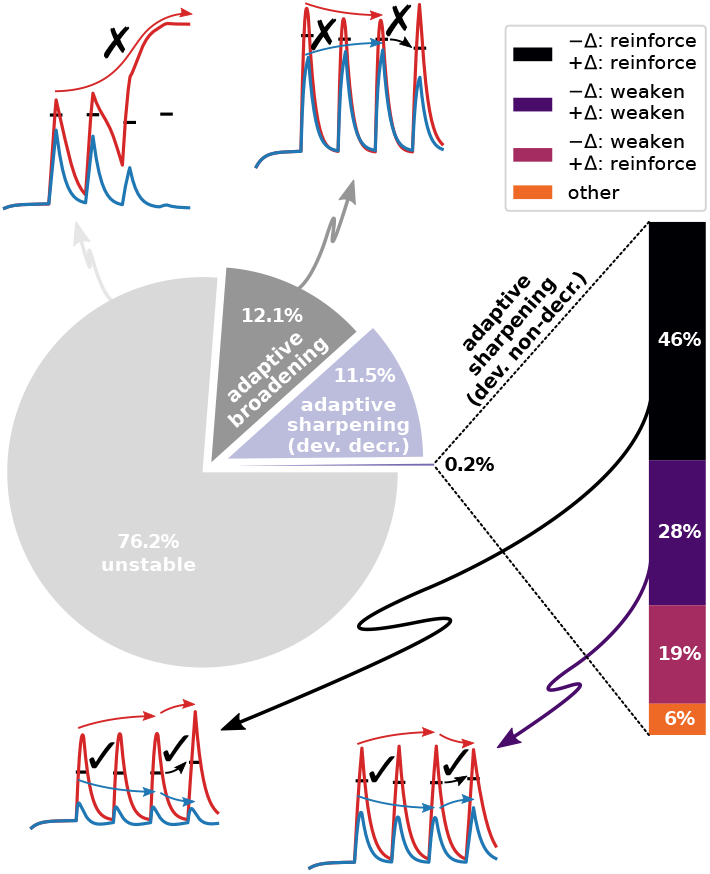
Deviance-driven complex tuning emerges innately from network connectivity configurations that facilitate adaptive sharpening and a deviance non-decreasing (DnD) response. Pie plot (left) shows the proportion, albeit small, of randomly sampled parameter configurations that produce all desired emergent phenomena (i.e., stable adaptive sharping of a neural representation’s tuning) or undesired emergent phenomena (i.e., unstable runaway excitation, adaptive broadening, and/or a deviance-driven decreasing response). Samples producing all desired phenomena are further categorized by the form of deviance-driven complex tuning they produce, whose relative proportions are presented in the barplot (right). Example simulations with unstable runaway excitation (upper left inset), adaptive broadening with a deviance-driven decreasing response (upper right inset), adaptive sharpening with a DnD response and deviance-driven complex tuning (bottom left and right insets).

One form of deviance-driven complex tuning was more robust than the others (responder rank reinforcement for both negative and positive deviants) and therefore emerged more prominently from the network despite parameter variability, occupying 46% of the valid parameter space. Despite this, the parameter-space responsible for producing the concurrent emergent phenomena of adaptive sharpening, deviance-driven complex tuning, and DnD responses was remarkably small. Through inspection of this underlying parameter-space, we were able to make a few inferences about the necessity of CI and SDI (Figure 4).

**Figure 4.**
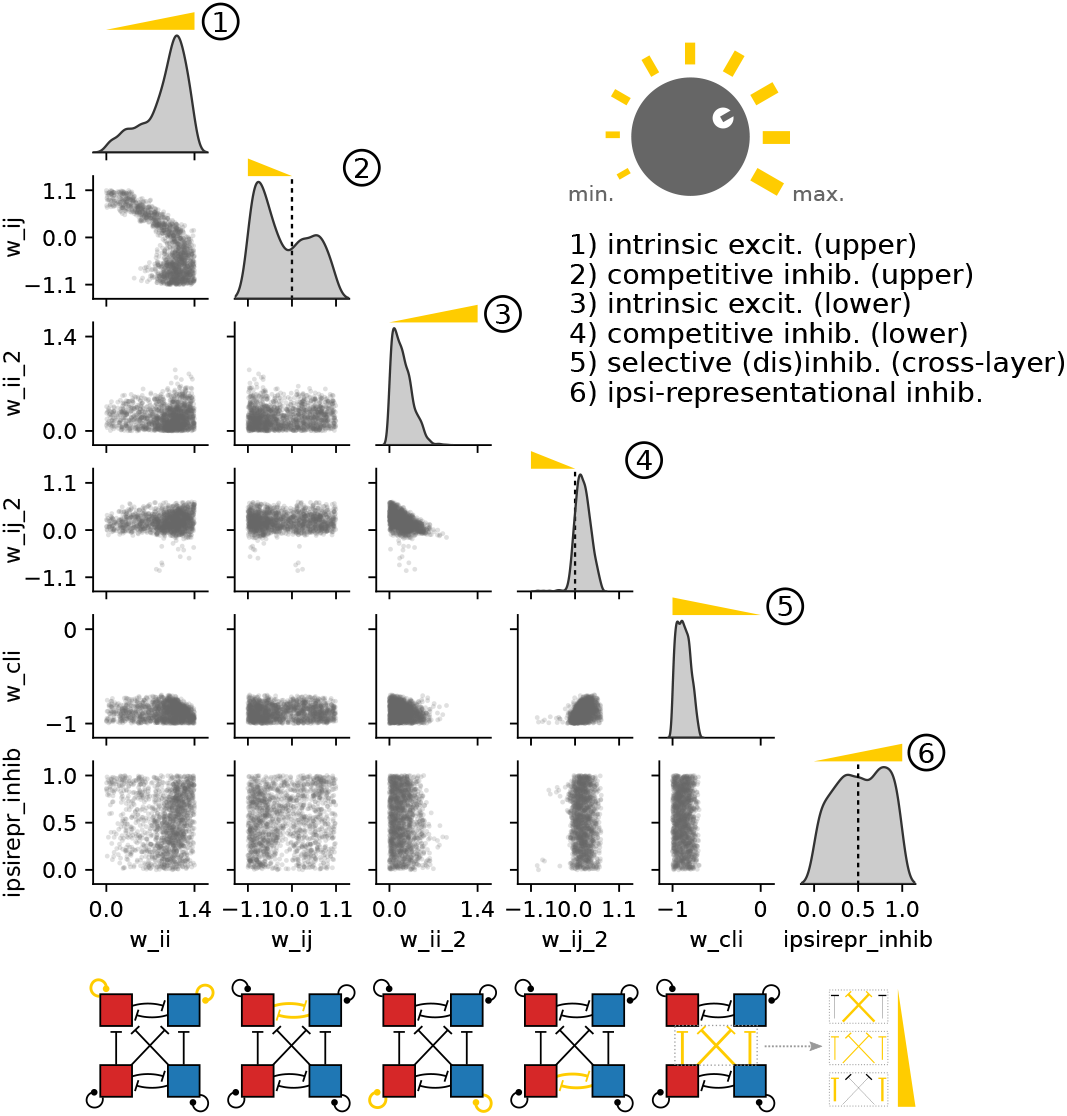
Adaptive sharpening, deviance-driven complex tuning, and the ability to produce deviance non-decreasing responses emerge from a small subset of the network connectivity parameter-space. Each point in a given scatter plot denotes a sample within the parameterspace that allowed the network to produce all desired emergent phenomena. Marginal kernel density estimates for each parameter are shown on the diagonal, the peaks of which demark the location of the maximum likelihood estimate for producing the desired emergent phenomena. Each of the six network parameters represents the strength of a pair of connections constrained by the relaxed CI+SDI motif (bottom). Yellow ramps (numbered 1-6) indicate the direction of increasing strength for each fundamental CI+SDI motif element mapped from a given underlying network parameter. For example, increased CI is induced by decreasing `w_ij` or `w_ij_2` starting at zero.

### Emergent deviance tuning dynamics are highly sensitive to network connectivity, but competitive inhibition promotes robustness

As graphically depicted at the bottom of Figure 4 (yellow connections), the loosened CI+SDI motif parameters comprise a positive (excitatory) recurrent connection weight for each of the upper layer units (`w_ii`; symmetric across units), a negative or positive connection weight between the upper layer units (`w_ij`) allowing for either CI or ME, two similar weight parameters for the lower layer (`w_ii_2` and `w_ij_2`), a negative (inhibitory) weight for the maximal cross-laminar inhibition (CLI) connection from the lower to upper layers that can have a maximal value of zero, and a [0, 1] scalar value that indicates the fraction of total CLI getting routed through connection weights between member units of the same neural representation (`ipsirepr_inhib`). Note that connectivity symmetry is enforced between the preferred and non-preferred representations at all times, allowing the CI+SDI motif to embody arbitrary anatomically-determined neural representations whose meaningful dynamics emerge from asymmetrical activation from thalamus or other brain area according to their stimulus, motor, or contextual mapping. Each randomly sampled parameter configuration that produced the aforementioned criteria of desired emergent phenomena is represented as a point in Figure 4. A kernel density estimate was used to approximate the marginal distribution for each parameter (Figure 4 diagonal subplots). Increased magnitudes of CI connections, along with other fundamental CI+SDI motif elements like SDI implemented through CLI, are indicated graphically for each parameter as a yellow ramp in Figure 4.

The first inference we made about the underlying network parameters that produce all three emergent phenomena (adaptive sharpening, deviance-driven complex tuning, and a DnD response) was that CI enhances robustness to parameter variability. The distribution of reciprocal connection weights in the upper layer (Figure 4, diagonal subplot 2) has its most prominent peak in the negative range of (`w_ij`) values, indicating that random parameter configurations with CI have a greater likelihood of producing the desired emergent phenomena than with ME. For the lower layer, ME was favorable over CI. The positive peak (i.e, `w_ij_2` *>* 0; Figure 4, diagonal subplot 4) indicates that the adaptive effects of CI are undesirable for the ensemble responses in this layer, likely due to the modulatory role the lower layer needs to have on the upper layer and that it needs to encode stimulus intensity rather than the adaptive influence of stimulus repetition. The robustness with which the lower layer encodes raw stimulus intensity allows it to better modulate the inhibitory landscape of the upper layer in response to a deviant shift in intensity.

Upon observing that CI increases the robustness of emergent deviance-tuning dynamics amidst connection variability, we characterized how sensitive such emergent dynamics were to time (specifically, the inter-stimulus interval [ISI]). Taking the connectivity parameter configurations that passed all criteria (i.e., exhibiting evoked responses with adaptive sharpening that were also DnD, as shown as point samples in Figure 4) for a 100 ms inter-stimulus interval (ISI), we simulated the network’s response with an extended or retracted ISI over a range of values from 50-200 ms to quantify the proportion of connectivity configurations that retained stability and DnD status (Figure 5). Connectivity configurations originally classified according to their emergent deviance tuning forms in Figure 3 fell into one of three major categories that each correspond to a unique region within parameter-space (Figure 5 inset with random parameter configurations plotted as points in 3-parameter dimensions): responder rank reinforcement for both negative and positive deviants (black), responder rank weakening for both negative and positive deviants (purple), and responder rank weakening (reinforcement) for negative (positive) deviants distinctively (pink). Of particular note, connectivity configurations with `w_ij` < 0 (i.e., CI instead of ME) predominantly led to responder rank reinforcement (black compared to pink and purple) and retained stability and DnD status across ISI values greater than 100 ms while other parameter configurations did not (Figure 5 solid and dotted traces).

**Figure 5.**
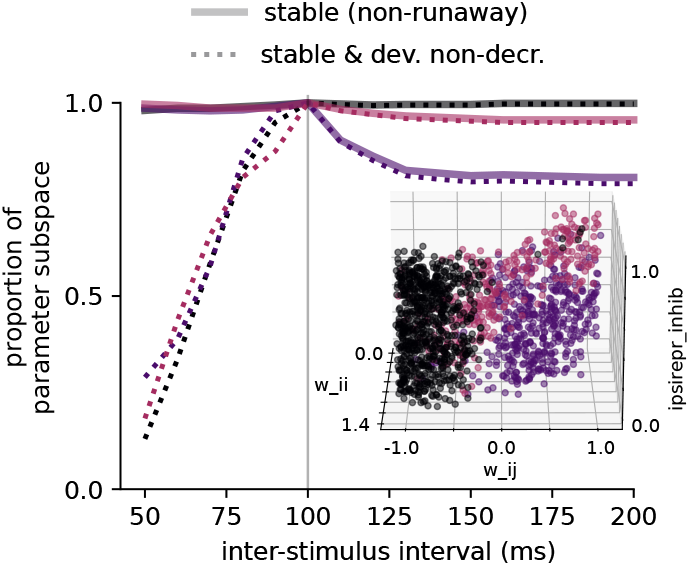
Deviance-driven dynamics are stable across a range of time intervals, particularly for networks with strong competitive inhibition (CI). Taking the connectivity parameter configurations sampled in Figure 4 that pass all criteria (i.e., exhibiting evoked responses with adaptive sharpening and are deviance non-decreasing [DnD]) for a 100 ms inter-stimulus interval (ISI), we extended (or retracted) the ISI over a range of values from 50-200 ms to quantify the proportion of connectivity configurations that retained stability and DnD status. Connectivity configurations originally classified according to their emergent deviance tuning dynamics (as indicated in Figure 3) fall into one of three major categories that each correspond to a unique region within parameter-space (inset): responder rank reinforcement for both negative and positive deviants (black), responder rank weakening for both negative and positive deviants (purple), and responder rank weakening (reinforcement) for negative (positive) deviants distinctively (pink). Of particular note, connectivity configurations with `w_ij` < 0 (i.e., producing CI instead of mutual excitation; black and pink) retained stability and DnD status across ISI values greater than 100 ms.

### Deviance-driven tuning requires strong selective (dis)inhibition

The second inference we made about the underlying network parameters that promote adaptive sharpening, deviance-driven complex tuning, and a DnD response was that strong SDI is crucial.

Across all valid parameter configurations, the distribution of CLI values peaks near `w_cli` = −1, at the lower end of its range (Figure 4). In fact, no valid parameter samples exist near `w_cli` = 0, indicating that having a network with SDI of maximal strength not only increases the likelihood of producing the desired emergent phenomena given a uniform random prior of other parameters, but is a requirement for producing the desired emergent phenomena at all. We emphasize that representing SDI as a uni-directional inhibitory connection between two layers in a network (i.e., CLI) is just one of many ways to implement SDI in a model that are functionally equivalent. The important thing is that inhibition must come from a separate group of neurons that share afferent input, which then modulates the E/I landscape of the group of neurons in which adaptive sharpening and deviance-driven complex tuning is measured.

### Inhibitory competition and selective cross-laminar (dis)inhibition regulate deviance tuning in a spiking neocortical column network

Up until this point, we explored network dynamics underlying deviance-driven complex tuning in a simplified spike rate model that approximates ensembles of neurons as neural mass units. While such a model conveniently simulates smooth time-varying spike rate functions that emerge from a network governed by only six parameters, we sought to test if our results extend to spiking dynamics in a biophyscially-detailed neocortical column network. Specifically, we found earlier in this study that CI increases the robustness of a network’s capacity for producing deviance-driven complex tuning and that SDI is critical for such phenomena to emerge at all: are these network mechanisms still relevant in a random, sparsely connected network that has been constrained with biophysical realism and anatomical connectivity rules? We also wished to account for the potential influence of thalamocortical adaption, a prominent component of stimulus-evoked responses in Neocortex absent from simulations in the neural mass model [27, 51, 66, 69, 70].

We specifically tested the hypothesis that at least one of the deviance-driven complex tuning scenarios of responder rank reinforcement, rank weakening, or rank reassignment (Figure 1d) can be produced within a biophysically-detailed neocortical column model constructed with CI between stimulus preferred and stimulus non-preferred ensembles and monosynaptic L6→L2/3 CLI. Further, we predicted that deviance-driven complex tuning would be disrupted by ablation of the L6→L2/3 inhibitory connection, a plausible anatomical pathway for SDI we built into the model. As elaborated on in Discussion, monosynaptic L6→L2/3 inhibition is implemented here as a hypothetical mechanism that warrants further experimental investigation. Our methods and results draw inspiration from prior work indicating a causal influence between stimulus-driven ensembles of L6 corticothalamic cells (CT) and the emergence of deviance-driven complex tuning in L2/3 [2], presumably through an established CLI pathway from L6 to the more superficial layers of the cortex [71, 72].

Using the Human Neocortical Neurosolver (HNN) modeling framework [55, 73], we simulated the process by which a neocortical column receives and processes repetitive and/or deviant stimulus-evoked exogenous perturbations from other brain regions such as the sensory thalamus (Figure 6). Constrained by prior experiments and neurophysiology data, the biophysically-detailed model contains neocortical layers 2/3, 5, and 6 and two distinct neuron ensembles in both L2/3 and L6 (preferred representation, red; non-preferred representation, blue) that receive different magnitudes of excitatory stimulus-evoked drive from exogenous sources (Figure 6a). Note that this model explicitly controls the activity of the granular layer (L4, the putative source of feedforward thalamic input) by defining its mean evoked spike rate rather than building L4 neurons into the model. L5 comprises only one functional ensemble. Inputs to the network come in the form of excitatory pre-synaptic spikes that arrive in either (1) L4 and L6 from lemniscal thalamus (proximal drive), or (2) L2/3 from non-lemniscal thalamus or high-order Neocortex (Figure 6b). Stimulus-evoked dynamics then propagate vertically (and laterally) throughout the network, governed by biophysical constraints within individual neurons and local network connections between neurons (Figure 6b). For baseline activity, the network was provided with excitatory Poisson drive (i.e., spike time events sampled from a Poisson distribution) to the somas of each neuron to establish sparsely firing activity that matches average rates from previous recordings in rodent barrel Neocortex (see Methods and Figure S2) [74, 75].

**Figure 6.**
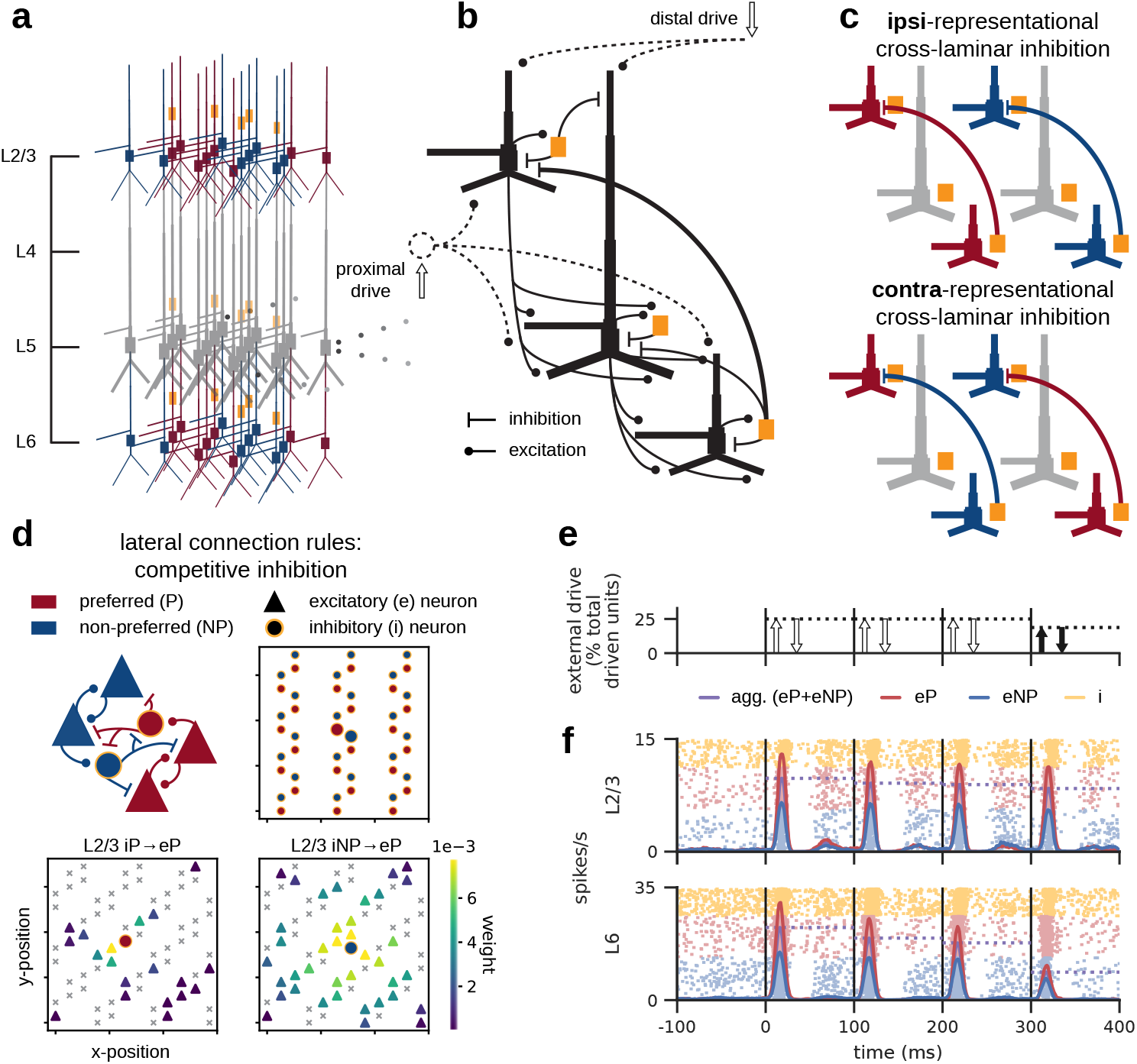
Competitive inhibition (CI) combined with cross-laminar selective (dis)inhibition can be extended to a spiking network model with biophysical realism and anatomical connectivity rules. (a) Overview of the model that contains neocortical layers 2/3, 5, and 6. L4 activity is not modeled, but rather controlled explicitly through proximal drive. Neurons in L2/3 and L6 are sorted into two distinct stimulus representations (preferred, red; non-preferred, blue) that make up functional ensembles and receive different magnitudes of excitatory stimulus-evoked drive from exogenous sources. (b) ‘Proximal’ drive (upward arrow) from lemniscal thalamus arrives at L4 and L6, while ‘distal’ drive (downward arrow) arrives at the uppermost extent of L2/3. Excitatory and inhibitory dynamics propagate throughout the network via local network connections. (c) L6→L2/3 cross-laminar inhibition (CLI) can have either ipsi-representational or contra-representational targets. Here, we constrained CLI to be 75% ipsi-representational. (d)Lateral connectivity between neurons within a given layer was parameterized for each neuron type using connection probability and distance-dependent maximal post-synaptic conductance. CI was implemented by increasing the probability and spatial conductance decay constant for cross-ensemble relative to within-ensemble inhibitory connections. For example, the connections from inhibitory interneurons to excitatory pyramidal neurons are more numerous and of greater strength between the red and blue representations than within a given representation. (e) The sequence of repetitive exogenous drives (proximal, then distal) that end with a negative amplitude deviant (solid black arrows). (f) Simulated aggregate spike raster and average spike rate in L2/3 and L6 across 40 random trials/networks.

SDI was implemented as L6→L2/3 CLI and can have either ipsi-representational or contra-representational targets (Figure 6c). For simplicity, we constrained CLI to be 75% ipsi-representational and 25% contra-representational; however, different configurations for this parameter could theoretically be used. Lateral connectivity rules between neurons were imposed by two parameters for each neuron type: connection probability and distance-dependent maximal post-synaptic conductance. CI was implemented by increasing the probability and spatial conductance decay constant for cross-ensemble inhibitory connections relative to cross-ensemble inhibitory connections (Figure 6d). For example, the connections from inhibitory interneurons to excitatory pyramidal neurons were more numerous and of greater strength between the preferred (red) and non-preferred (blue) representations than within a given representation.

A single simulated evoked response trial was composed of a sequence of exogenous drives that represents the afferent and contextual exogenous inputs received from other brain areas evoked by four repetitive stimuli administered 100 ms apart. Building from prior work [55, 76–78], each repetition contained a proximal and distal drive at ∼12 ms and ∼35 ms post-stimulus, respectively, that were slightly weaker or stronger for the deviant (fourth) repetition (Figure 6e). As described in more detail in Methods, evoked drives were implemented as pre-synaptic spike events sampled at times from a Gaussian distribution, entering the cortex at a specific location (proximal or distal) and targeting specific post-synaptic cells. Evoked drive strength was modulated for deviant afferent drive by decreasing or increasing the number of total driven cells from a baseline value for both proximal and distal drives. To account for the effect of depressing thalamocortical adaptation [64, 66, 70], the synaptic strength (i.e., maximal postsynaptic conductance) of proximal drives attenuate at a rate of 8% per repetition. Importantly, the ratio of proximal drive strength between the two ensembles remained constant across the entire sequence of evoked drives. Here, we focused on the negative deviant case (i.e., a decrease in drive strength for the fourth repetition) because network dynamics that refrain from attenuating despite decreasing excitatory input are among the most challenging phenomena to imbue in a spiking neural network, particularly when simple disinhibition isn’t at play.

Upon building the model with symmetrical CI between ensembles of neurons in L2/3 and L6 and added L6→L2/3 CLI that, some preliminary parameter tuning was needed prior to testing if the network could produce deviance-driven complex tuning in L2/3. Specifically, we tuned the model to produce sparse baseline firing rates and non-decreasing positive *and* negative deviance-evoked responses (averaged across all neurons in L2/3, stimulus-preferred and non-preferred), as outlined in detail in Methods. In a representative example batch simulation (consisting of 40 random trials) of the final tuned model (Figure 6e,f), the number of cells targeted by the evoked drive (empty arrows; Figure 6e) were contained to be constant across each stimulus repetition until the end, at which point a deviant decrease (−Δ) occurs (solid black arrows). Note that we designed deviant afferent drives to maintain the same ratio of post-synaptic targets between ensembles (preferred to non-preferred) as for the non-deviant drives, ensuring that any emergent shifts induced by deviance were the product of intrinsic network dynamics rather than an external shift in, e.g., thalamic tuning. The mean single-unit spike rate across simulation trials and neurons increased following each proximal and distal drive in L2/3 (red and blue traces), but only for proximal drive in L6 due to the fact that L6 lacks direct distal input (Figure 6f). For both L2/3 and L6, the mean evoked response in the preferred ensemble (red) always peaked higher than for the non-preferred ensemble (blue), reflecting the difference in drive strength received by each ensemble. The mean peak response across ensembles (horizontal purple dotted line) slightly decreases with each repetition due too the modeled effect of thalamocortical adaption that decreases the synaptic strength of afferent drive with each stimulus repetition. On the final repetition of proximal and distal afferent drives, mean L6 peak activity decreases significantly with the deviant change in proximal drive, but L2/3 evoked activity merely keeps pace with thalamocortical adaptation (Figure 7a, purple).

**Figure 7.**
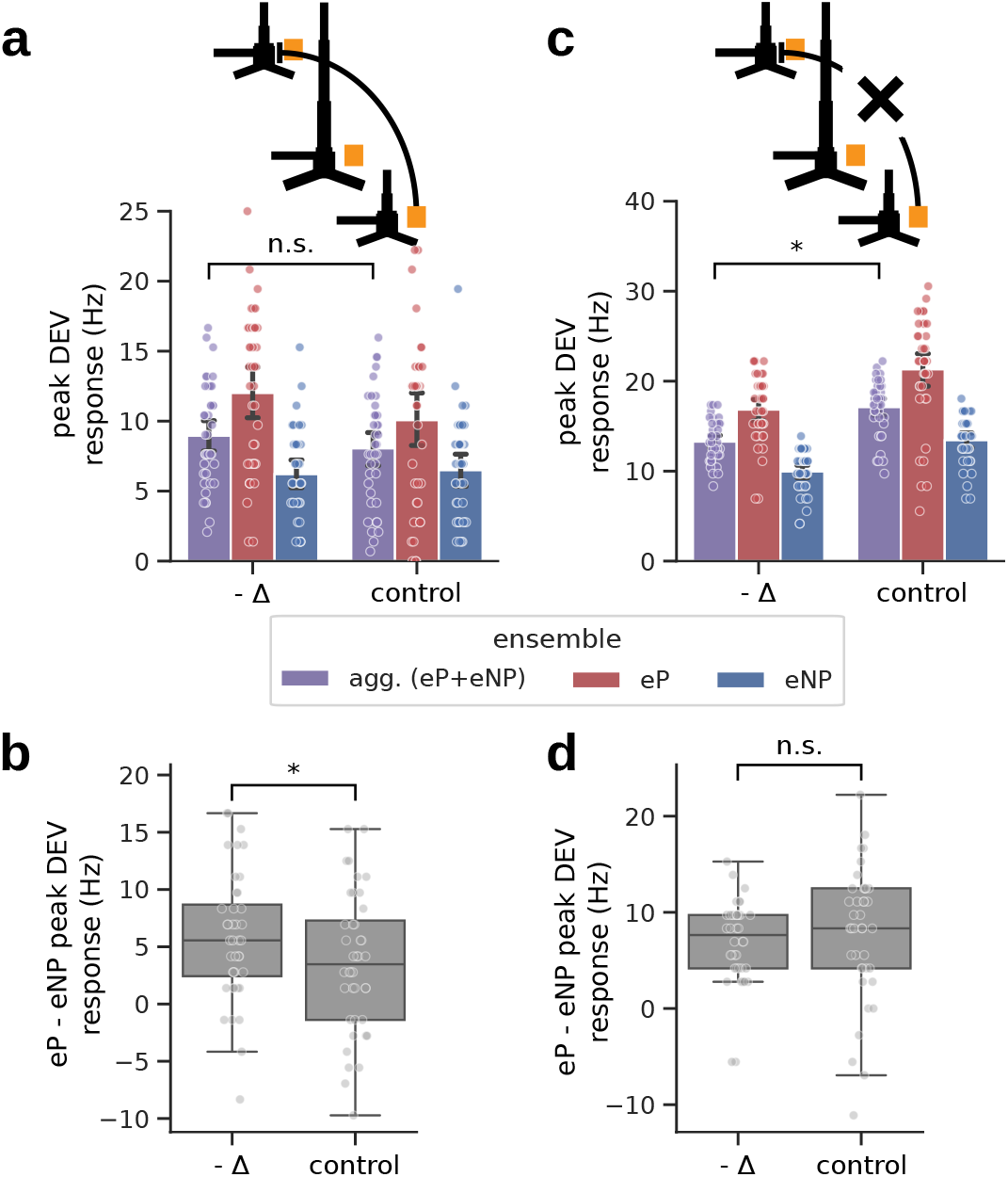
Despite the added constraints of biophysical realism and anatomical connectivity rules, deviance-driven complex tuning emerges with competitive inhibition and cross-laminar selective (dis)inhibition (SDI) in a spiking neocortical column model. Consistent with dynamics of the neural mass model, SDI is essential. (a) Mean deviant evoked response across n=40 random trials/networks with 95% confidence interval estimated via bootstrap over 1000 permutations. For a negative deviant (−Δ) where the afferent drive to the network decreases, the average neural response aggregated across all excitatory neurons in L2/3 (purple) doesn’t decrease compared to the non-deviant control. (b) Box plot showing the quartiles of the difference between preferred (eP) and non-preferred (nP) evoked responses across paired samples [red vs. blue in (a)], each pair from a given random network. The difference in evoked response is larger for the negative deviant compared to control indicating significant deviance-driven tuning, specifically, responder rank reinforcement. (c) With ablation of the cross-laminar SDI connection, the decrease in afferent drive correspondingly decreases the mean deviant evoked response across all ensembles compared to control. (d) Similar to (c), except where ablation of the cross-laminar SDI connection correspondingly removes deviance-driven tuning. ^*^P-value< 0.05.

After tuning the model, we tested for the presence of deviance-driven complex tuning in L2/3 by comparing the difference between preferred (eP) and non-preferred (nP) evoked peak responses on deviant versus repetition-matched non-deviant (control) trials (Figure 7a,b). For a negative deviant where the afferent drive to the network decreases, the average neural response aggregated across all excitatory neurons in L2/3 (purple) did not significantly differ from that of the non-deviant control (Figure 7a; two-tailed Wilcoxon signed-rank test across trials/networks, P-value=0.217). When grouped according to their respective ensembles (red vs. blue), however, the response of excitatory neurons in the preferred ensemble (eP) were visibly higher than that of the non-preferred ensemble (eNP) for both the deviant and control, as expected given differing levels of evoked drive. Deviance-driven complex tuning emerged in the model as an increased difference between the eP and eNP peak responses (i.e., responder rank reinforcement; Figure 7b; upper-tailed Wilcoxon signed-rank test, P-value=0.011).

Further, we verified that the emergence of deviance-driven complex tuning was sensitive to removal of SDI by ablating the CLI connection in the model (Figure 7b,c). With ablation of the CLI connection from L6 to L2/3 (Figure 7c), the decrease in afferent drive correspondingly decreases the aggregate deviant evoked response compared to control (P-value=4.19 × 10^−7^), demonstrating that a fundamental condition of DD (maintaining, if not increasing, the net salience of the neocortical area’s output) was lost. Significant deviance-driven complex tuning was also lost (Figure 7d; P-value=0.916). Consistent with our theoretic results from the simple neural mass model earlier in this study, CLI ablation didn’t significantly change baseline activity in L2/3 (data not shown), but rather, shifted how the two different ensembles responded to stimulus-evoked drive and each other.

It’s important to note that simulations with this particular tuning of the biophysically-detailed model do not rule out the possibility that the other forms of deviance-driven complex tuning (i.e., responder rank weakening and rank reassignment as depicted in Figure 1d) may indeed be achievable with different network configurations (e.g., with a different balance of ipsi- versus contra-representational CLI or with a different level of within-ensemble excitatory synaptic strength). Here, we demonstrate that despite the added constraints of biophysical realism and anatomical connectivity rules, deviance-driven complex tuning emerges with CI and cross-laminar SDI in a spiking neocortical column model. Consistent with dynamics of the neural mass model and in support of our overarching hypothesis, SDI is essential.

## Discussion

In this study, we introduce and explore a novel theory that the neural dynamics subserving DD in the early-latency phase of a stimulus-evoked response are the product of competing neural representations, each representation encoding specific features of the stimulus and/or contextual environment. The network mechanisms of this process we term CI and SDI. Together, CI and SDI imply a specialized network connectivity motif that promotes a handful of emergent neural phenomena previously unaccounted for by theoretical models of predictive processing within the Neocortex.

CI is the process by which at least two neural representations embodied by distinct (but not necessarily mutually exclusive) ensembles of neurons reciprocally inhibit each other and augment the difference in stimulus tuning with each repetition of the same stimulus. SDI is the process by which evoked afferent drive modulates the speed and extent of CI via feed-forward inhibition, establishing a novel E/I landscape for the competing ensembles to navigate whenever the raw intensity of a stimulus (or its relative composition of features) deviate. We first showed that deviance-driven complex tuning emerges with CI and SDI in a 4-dimensional neural mass model, demonstrating sufficiency of the underlying CI+SDI motif for facilitating the most fundamental criteria of DD: selective amplification of the deviance-driven response from a subset of neurons at the expense of others without decreasing the net output of the network.

Receptive fields shift from one context to the next based on stimulus history [51, 60], the mechanisms for which have typically been associated with top-down or bottom-up modulation from other brain regions relative to the neocortical hierarchy (e.g., as in hierarchical predictive coding [5, 7, 79, 80]). Inspired by prior work that demonstrates fast time-scale inhibition can dynamically shape functional neural representations before and after network learning [42, 45, 46, 58, 75, 81–83], we establish that recurrent CI allows for recent historical context (i.e., the basis of a sensory prediction) to be computed and stored locally through a stimulus’ representational evolution in state space, as induced by repetitive afferent drive. This mechanism is analogous to that of the synaptic theory of working memory whereby slow dynamics of residual pre-synaptic calcium shape recurrent neural network representations of items stored in working memory, except that here we propose that Sensory Neocortex leverages post-synaptic activity (which can lead to DD encoded in spike rates as an emergent property) to store short-term memory instead of pre-synaptic activity [84, 85]. The synaptic theory of working memory relies on calcium-mediated synaptic short-term facilitation, a phenomenon common in high-order neocortical areas like the Prefrontal Neocortex yet significantly less common in primary sensory areas where synapses are largely depressing [86].

CI relies on two features of functional cell-to-cell connectivity: 1) strong recurrent connections among excitatory neurons of similar stimulus preference that 2) innervate fast-spiking PV interneurons with post-synaptic targets of a different stimulus preference. Both conditions receive strong support from in vivo calcium imaging and in vitro slice recordings in rodent visual Neocortex [39–42] and are further supported by anatomical reconstruction via electron microscopy [87]. However, the CI+SDI motify need not rely solely on a static anatomical connectome. Functional PV organization may shift dynamically to enforce CI when under the influence of other interneuron cell types common to L2/3, such as vasoactive intestinal peptide-expressing (VIP) and somatostatin-expressing (SST) interneurons that are known to modulate functional connectivity and play a role in encoding novel stimuli [88]. VIP in particular have been shown to interact with SST to help regulate the gain of neural circuits in service of enhanced stimulus specificity without compromising network stability [29]. Placing the CI+SDI motif within the context of ensembles rather than single neurons promotes robustness despite variance in single neuron activation notable in the sparse stimulus-driven representations of Sensory Neocortex [75].

The results of the current study demonstrate that CI can prime the network by facilitating adapting neural representations (specifically, adaptive sharpening), a previously established phenomenon of neural tuning that had until now lacked theoretical backing for shared circuitry with DD [51]. Such alignment is congruent with the broad observation that over longer time-scales, perceptual experience facilitates increased specificity (i.e., sparseness) in a stimulus’ neural representation [89]. We further establish that while SDI is necessary for deviance-driven complex tuning, CI is not. Rather, CI enhances robustness to connectivity and temporal variation for the co-emergence of adaptive sharpening and deviance-driven complex tuning. This latter result is likely due to the existence of a highly sensitive parameter regime where intrinsic differences in afferent excitation are able to provide enough repetition-driven adaptation that SDI can still instigate a deviance-driven shift in tuning. While this matter requires further investigation, our cumulative results indicate that the CI+SDI motif is at minimum necessary for the robustness of deviance-driven shifts in neural tuning given intrinsic variance contained in most (if not all) real-life neural circuits and processes.

The CI+SDI motif’s ability to produce deviance-driven complex tuning are conserved across levels of physiological abstraction. In the final part of this study, we extended the CI+SDI motif from a reduced network with hypothetical neural mass units producing firing rates to a spiking neocortical column network constrained with biophysical realism and anatomical connectivity rules (Figure 6). Our results validate that deviance-driven complex tuning can emerge from spiking interactions in and between ensembles of distinct neurons and that such tuning is particularly sensitive to SDI, as expected from results obtained using the neural mass model (Figure 7). While we introduced assumptions about the neocortical elements underlying our proposed CI+SDI motif, these assumptions in no way compromise the functional significance of our theory. In particular, we assumed that SDI is mediated by monosynaptic L6→L2/3 CLI. While there is experimental evidence for a causal link between specific ensembles of stimulus-driven L6 CT and the emergence of deviance-driven complex tuning [2], as well as an established cross-laminar interneuron in L6 that provides inhibitory gain modulation onto the more superficial layers of the neorcortex [71, 72], L6 might only influence L2/3 through polysynaptic connections.

L6 is compelling as a potential source of SDI for a number of other reasons given the unique stimulus tuning properties of L6 CT and their anatomical connectivity within thalamocortical circuitry. L6 CT exhibit distinctly sharp tuning profiles to external stimuli [90], allowing them to encode specific features of the sensory or contextual environment in a robust manner as sparse ensembles. They also receive direct projections from lemniscal thalamus and send direct projections to both lemniscal and non-lemniscal/high-order thalamus, giving CT a crucial role in mediating thalamocortical feedback and modulating contextual input entering various layers of Neocortex [90–94].

DD is typically associated with a pronounced increase in evoked spike rates across neurons of an experimentally measured population [11–13]. Here, we focused on early-latency (< 100 ms) evoked response phenomena associated with DD that involves the selective amplification of the response of some neurons over others (i.e., as exemplified by deviance-driven complex tuning; Figure 1c,d), especially highlighting the case where the mean response across neurons remains non-decreasing despite a deviant decrease in afferent drive. Our hypothesized mechanisms of CI modulated by SDI intrinsically produce ensemble-level evoked responses that shift over time and space/neurons, implying that the conjunctive (i.e., aggregate) representation across ensembles transiently shifts as well. From the perspective of an observer who is naive to the complete anatomical connectivity of a random sample of neurons and the ensembles they belong to (as is the case in most electrophysiology experiments), the early-latency deviant response is predicted to correspond to a notable shift in stimulus decoding accuracy relative to baseline. The shift in stimulus encoding can either increase or decrease depending on the specific form of deviance-driven complex tuning: responder rank reinforcement corresponds to increased decoding accuracy, whereas responder rank weakening or reassignment corresponds to decreased decoding accuracy. Based on the results of this study, responder rank reinforcement is the most robust form of deviance-driven complex tuning and emerges exclusively from networks with CI and strong SDI. Future in vivo experiments can test this theoretical prediction, along with whether functional ensembles of neurons classified according to their baseline tuning profile shift on deviant trials and whether that shift critically depends on cross-ensemble inhibition defined in the CI+SDI motif.

## Methods

### Neural mass model simulations

The Wilson-Cowan neural mass model given by Equation 1 contained up to four units with a sigmoid activation function of the form

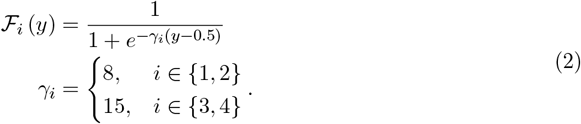

The response of each unit thus had a dynamic range in (0, 1), 0 representing a quiescent ensemble (i.e., with a spike rate of 0), 1 representing a maximally active ensemble (i.e., with a saturated peak spike rate). The steepness parameter *γ* was symmetrical across the stimulus-preferred and non-preferred representations and fixed at a smaller value for units in the upper layer (units 1 and 2) so that the dynamic ranges of their activation functions were greater, or slower acting, than for units of the lower layer (units 3 and 4). The time-varying injected current parameter *I*(*t*) took a value of 0.01 at baseline and 0.55 and 0.45 for a duration of 20 ms during evoked drive for the preferred and non-preferred units, respectively. The magnitude of evoked drive increased or decreased by a multiple of 0.2 for positive or negative deviants, respectively. We simulated network dynamics using the Runge-Kutta method (RK4) with integration time step Δ*t* = 0.1 ms, time constant *τ* = 20 ms, and initial conditions *x*_*i*_ = 0 ∀ *i* ∈ {1, 2, 3, 4} (Equation 1).

For the initial proof-of-principle of emergent dynamics examined further in this study, the specific connection weights for the model simulated in Figure 2c,d were selected using an optimization routine that targeted the mean deviance-driven response across units in the upper layer. Pairs of connection weights constrained by symmetry between the stimulus-preferred and non-preferred representations, as defined in the fundamental CI+SDI motif, were allowed to vary during optimization, but `ipsirepr_inhib` was held fixed at 0.75. We first set parameter bounds that limited runaway excitation/saturation across units (`w_ii` ∈ (0.1, 1), `w_ij` ∈ (−1.1, −0.2), `w_ii_2` ∈ (0.1, 1), `w_ij_2` ∈ (−1.1, −0.2), and `w_cli` ∈ (−1, −0.1), see Figure 4 for a graphical depiction of these parameters) and then ran a gradient decent optimization routine (SciPy’s L-BFGS-B implementation) over those ranges to minimize loss ℒ given by

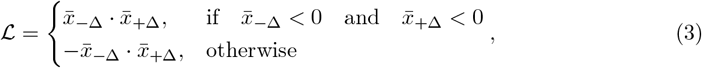

where 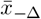 and 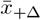 denote the mean peak firing rates across units in the upper layer (*x*_1_ and *x*_2_) for the negative and positive evoked responses, respectively, minus that of the non-deviant evoked response. Note that ℒ in Equation 3 is greater than zero iff either 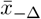 or 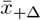 fall below 0, allowing the optimization routine to minimize the extent a deviant evoked response decreases relative to a non-deviant response. The 2-dimensional model simulation of CI without SDI shown in Figure 2b uses the same optimized parameters as that in Figure 2b, except with the upper and lower units disconnected.

While our goal was to observe if/how deviance-driven complex tuning might emerge from the model, it’s important to note that we never explicitly forced the model to produce any of the specific forms of deviance-driven complex tuning (responder rank reinforcement, weakening, or reassignment, as depicted in Figure 1d). Instead, we optimized the model to produce a mean evoked firing rate that increases (or at least does not decrease) with deviant afferent drive regardless of the direction of the deviant change in stimulus amplitude, a fundamental aspect of DD and constraint on emergent dynamics. We had no prior guarantees that a non-decreasing mean deviance-driven response across ensemble representations could be achieved by this system given our CI+SDI network motif; however, we reasoned that if the average rate didn’t decrease on a deviant, the underlying system would be forced to compensate by amplifying the response of one, and only one, of the constituent units in the upper layer. If a positive deviant were to increase the evoked response of both units, the system would be overwhelmingly governed by afferent excitation and result in a decrease in the mean evoked response on a negative deviant. If, on the other hand, a positive deviant were to decrease the evoked response of both units, the system would be overwhelmingly governed by cross-laminar inhibition and result in an increase in the the mean evoked response on a negative deviant (i.e., from acute disinhibition).

Source code for running simulations and optimization of the neural mass model is available at https://github.com/rythorpe/dev_detect_wc.

### Biophysically-detailed model simulations

All simulations of the biophysically-detaild model were run using HNN-core [73], a modular and expandable Python implementation of HNN [55]. HNN is based on a pre-constructed and pre-tuned model of a laminated neocortical circuit under thalamocortical and cortico-cortical influences. While creation of HNN was originally inspired by the desire to interpret the cellular and microcircuit origin of human EEG/MEG signals by relating the primary electrical currents (i.e., current source dipoles) in pyramidal neuron dendrites to a variety of commonly measured neurophysiology data types including local network spiking activity, LFP, and laminar current-source density (CSD), it was built using canonical features of neocortical dynamics and circuitry conserved across rodent, non-human primate, and human anatomy, thus allowing cross-species inference [55, 76, 77]. Here we used the HNN model to validate our theoretical results obtained in a neural mass model of ensemble spike rates to spiking dynamics between various neuron types and layers as constrained by the anatomy of a neocortical column.

Aside from stochastic baseline drive that is provided as 20 Hz Poisson pre-synaptic spike events with a 300 ms burn-in, activation of the HNN network is achieved by defining patterns of action potentials representing activity in exogenous brain areas (e.g., thalamus or high-order Neocortex) that drive the local network through layer specific patterns of excitatory synaptic inputs. Synaptic inputs that drive the network target the dendrites of excitatory pyramidal cells and the somas of inhibitory basket cells. One pathway of input simulates perturbations ascending from the peripheral nervous system, through the lemniscal thalamus to the granular layer (L4), and finally propagating to the proximal dendrites of the pyramidal neurons in L2/3 and L5, as well as the distal dendrites of pyramidal neurons in L6 [95]. These inputs are referred to as proximal drive. The other pathway of input comes from the non-lemniscal thalamus or high-order Neocortex and synapses in L2/3, targeting the distal dendrites of the pyramidal neurons. These inputs are referred to as distal drive [55].

It is important to acknowledge that there are numerous types of neocortical column models with varying levels of detail; however, this model was selected for this study because it balances biophysical detail with simplifying assumptions that make manipulations of within-layer and between-layer connectivity tractable and simulations computationally efficient compared to other large-scale models. Namely, there are a handful of cell types, each composed of up to a nine compartments governed by Hodgkin-Huxley cable equations implemented on the backend in NEURON [96]. By default, the HNN model has two primary layers (L2/3 and L5), however, we added L6 (comprised of two cell types, pyramidal cells and inhibitory basket cells) for the present study. Neurons within a given layer were placed at regular intervals in the laminar plane such that we could modulate the degree of lateral and vertical connectivity through three parameters: maximal synaptic conductance, horizontal spatial decay (i.e., the rate at which the synaptic conductance decays with horizontal distance between two neurons), and connection probability (see Figure 6 for a graphical depiction of the network connectivity rules). The position of neurons belonging to the preferred (P) and non-preferred (NP) representations were interleaved across a hexagonal 2D mesh within each layer to ensure that excitation within and CI between ensembles was symmetrical despite the 1:3 proportion of inhibitory-to-excitatory neurons.

Consistent with prior recordings that observed that L4 and L6 neurons receive direct afferent projections from sensory thalamus [97] and that stimulus-driven activation of local inhibition generally proceeds excitation in distinct representational ensembles [75, 98], synaptic delays of proximal drive in our model were chosen to activate L6 apical dendrites first, followed by the activation of L2/3 inhibitory interneurons and then excitatory pyramidal neurons. Stimulus-driven activity in L2/3 was therefore shaped primarily by the strength of proximal drive propagating from L4 combined with polysynaptic activity routed through L6. Given the purpose of this model and the scope of phenomena we wished to observe, L4 and L5 were assumed to primarily act as relay nodes within the column. This assumption does not diminish the fact that L4 and L5 are known to play complex and crucial roles in other types of cortico-cortical and thalamocortical processing, such as in in the dynamic shaping of spatial receptive fields in L4 neurons [27, 43, 99] and dendritic integration in L5 pyramidal cells [100, 101].

### Biophysically-detailed model parameter tuning

Tuning of the biophysically-detailed model involved an iterative procedure until the target baseline (Figure S2) and evoked spike rates were achieved. We modified the network connection weights and the rate of thalamocortical adaptation within the constraints of symmetrical CI and SDI via L6→L2/3 CLI from the base values distributed with HNN-core [73] to prevent excessive correlations across neurons so that baseline and evoked spike rates remained sensitive to stimulus history [102]. Simulating the response of a particular network configuration consisted of running a batch across 40 random instantiations of the final tuned model, for which aggregate spiking activity is shown in Figure 6f, with random input spike times (Poisson-distributed for baseline drive, Gaussian-distributed for evoked) and random network connections (uniformly sampled among neuron pairs based on an average probability defined for cell type). Due to the highly non-linear and cost-intensive nature of running simulations in this model, all tuning was conducted manually to produce resting-state spiking activity consistent with in vivo mouse recordings from prior studies [74, 75]. The procedure was as follows.

1. Fix cell positions, morphology and biophysics.
2. Connect Poisson drive to the soma of each cell independently.
3. In the disconnected network, tune Poisson drive synaptic weight parameters to achieve the disconnected (intermediate) average baseline firing rate for each cell type, which is a fraction of the final target average firing rate for each cell type.
4. With the Poisson drive synaptic weight parameters fixed, provisionally fix the local network connectivity parameters (this includes a pairwise connection probability for each connection type and spatial synaptic weight decay).
5. Instantiate the random network connections with a seeded psuedorandom number generator.
6. Tune maximal conductance (synaptic strength) parameters for each connection type in the local network:
  6.1 Achieve the target connected (final) average baseline firing rate for each cell type [74, 75].
  6.2 Achieve limited spatiotemporal spike correlations by visual inspection of 2 seconds of simulated baseline activity.
  6.2 If the network cannot accommodate these achievements with a manual parameter search, return to (4).
7. With local network connectivity parameters provisionally fixed, tune maximal conductance (synaptic strength) and thalamocortical adaptation parameters for each type of evoked drive (proximal and distal):
  7.1 Achieve reasonable evoked spike rates that don’t decrease their peak magnitude (averaged across ensembles) for both positive *and* negative deviants.
  7.2 Achieve non-saturating evoked spike rates in each neural population.
  7.3 Achieve evoked spike rates that appear to be sensitive to prior evoked activity.
  7.4 If the network cannot accommodate these achievements with a manual parameter search, return to (6).

The final tuned parameters and source code for running simulations of the biophysically-detailed model are available at https://github.com/rythorpe/hnn-core/tree/L6_model.

## Acknowledgments

We thank Scott Susi, Christopher Deister, and Jakob Voigts for helpful discussion on possible circuit motifs underlying deviant sensory processing, neocortical and thalamocortical anatomy, realistic spiking behavior in selected cell types, as well as the potential limitations of simplifying assumptions in the biophysically-detailed model used in this study. We also thank Nicholas Tolley for guidance on implementing parameter sweeps via high-performance computing and providing insight and discussion regarding avenues for statistical inference. Model simulations were conducted using computational resources and services at the Center for Computation and Visualization, Brown University. This study was supported by the National Institute of Neurological Disorders and Stroke (R01NS108414 and U24NS129945), and the National Institute of Mental Health (T32MH115895 and P50MH109429) of the National Institutes of Health. The content is solely the responsibility of the authors and does not necessarily represent the official views of the National Institutes of Health.

## Supplementary Material

**Figure S1:**
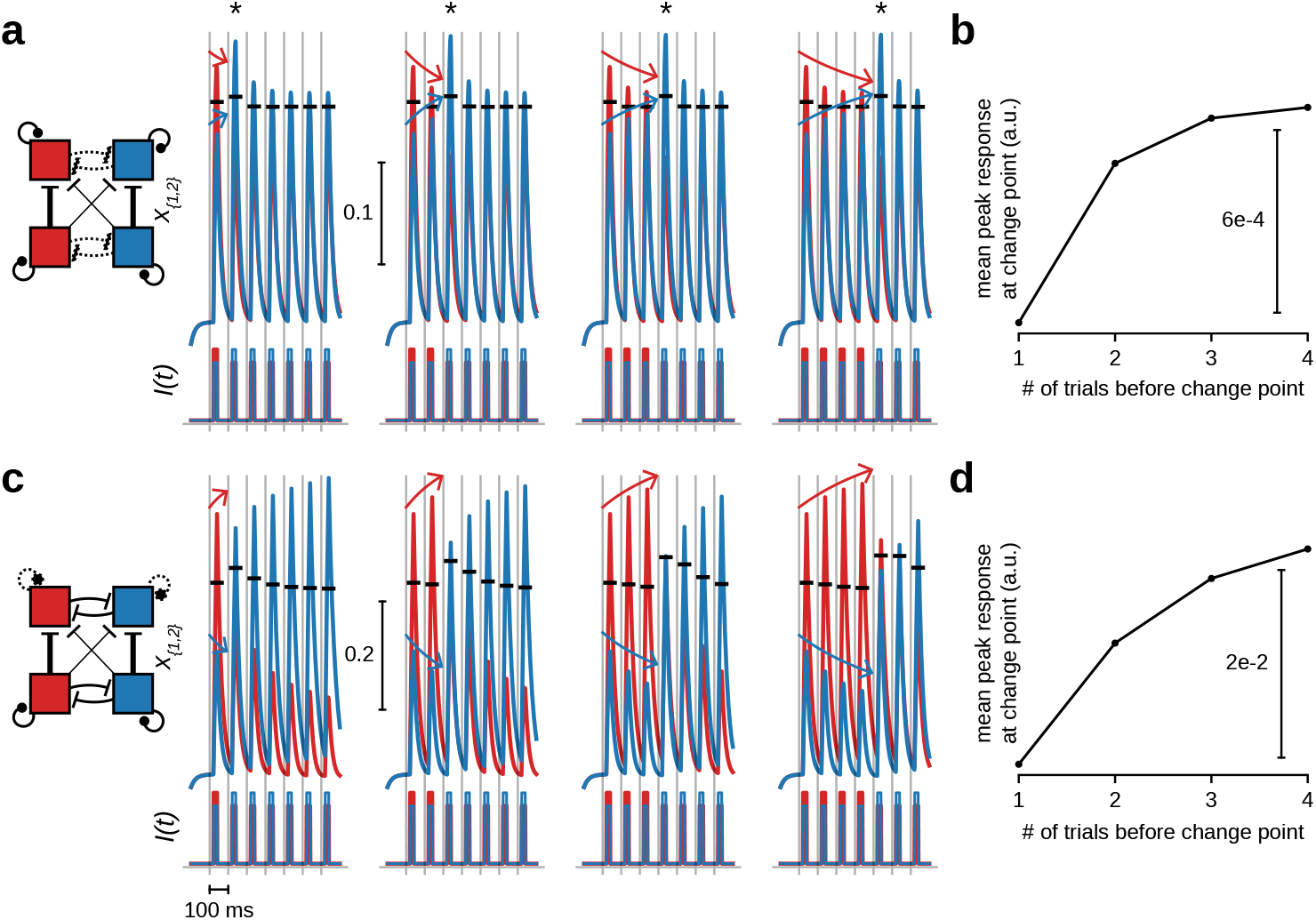
Competitive inhibition within the upper layer leads to ensemble priming, a network mechanism for context storage and history dependence. With pronounced within-layer inhibition (a) or within-unit excitation (c) that causes evoked responses between the upper layer units (i.e., *x*_1_ and *x*_2_) to either converge (adaptive broadening) or diverge (adaptive sharpening) with more stimulus repetitions of a particular feature preference, the average deviance-induced evoked response on a change point (indicated by an asterisk) increases (b,d). Note that in this stimulus paradigm, the relative balance of salient stimulus (or contextual) features serving as inputs to the system can be repeated and stored over time (e.g., red stimulus component *>* blue stimulus component or vice versa), as indicated by the history-dependent shift in deviant response magnitude.

**Figure S2:**
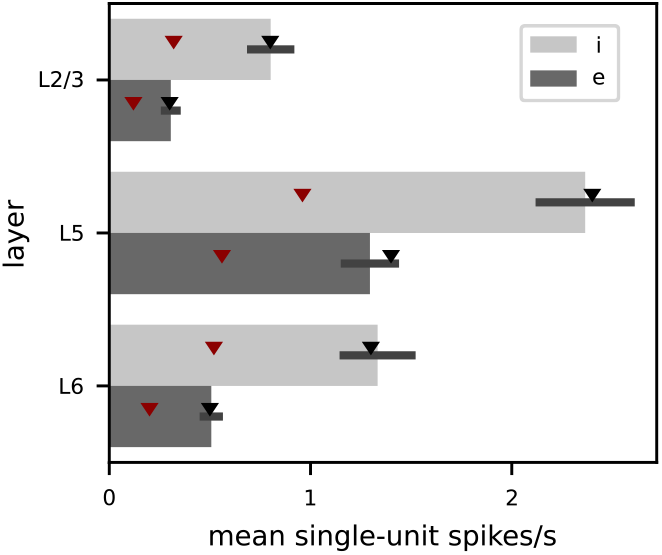
Target (triangles) and achieved (bars) mean baseline spike rates used for tuning the baseline activity of the biophyscially-detailed model. Poisson drive parameters were first tuned to match a fraction (0.4, red) of the final target baseline spike rates (black) in a disconnected network. Then the network connectivity was instantiated and tuned to increase the mean baseline spike rates from their intermediate to final target values (from top to bottom, L2/3-L6, 0.8, 0.3, 2.4, 1.4, 1.3, and 0.5 Hz). Tuning was considered complete once the final target fell within one standard deviation of the simulated mean (S.E., black error bars).

## Notes

### Competing Interest Statement

The authors have declared no competing interest.

### Summary of Updates

Introduction updated to clarify motivation and framing; Figure 3 split into two separate figures (now Figure 3 and Figure 4) with accompanying text in Results to streamline flow of logic.

